# Senescent astrocytic deposits drive cognitive decline by disrupting tripartite synapse in the aging brain

**DOI:** 10.1101/2025.10.31.685958

**Authors:** Er-Jin Wang, Chen Ming, Yi-Ting Wang, Shi-Jia Wang, Zi-Han Song, Kai Yang, Jin-Tao Li, Xu-Xu Zhuang, Wei Wang, Ming-Yue Wu, Li-Ming Xie, Zhengyu Ren, Chun-Ping Liu, Lei Wang, Yong-Gang Yao, Min Li, King-Ho Cheung, Aston Jiaxi Wu, Han-Ming Shen, Huanxing Su, Zhenyu Yue, Jia-Hong Lu

**Author notes:** These authors contributed equally to this work as the first authors.

## Abstract

Brain aging involves synapse decline, with astrocytes playing a key role in synapse homeostasis. However, the impact of astrocyte senescence on synaptic dysfunction and cognitive decline remains unclear. Here, we identified a hallmark of aging astrocytes—Senescent Astrocytic Deposits (SAD) observed at aged rodents, macaques, and human hippocampal astrocytic processes —that is associated with tripartite synapse dysfunction and memory decline. Laser capture microdissection-coupled mass spectrometry (LCM-MS), spatial transcriptome analysis and 3D electron microscopy revealed that SAD are abnormal protein deposits at the processes of ApoE-high expression astrocyte subtype and associated with dysfunctional tripartite synapses. Using a transgenic mouse (*Nrbf2* knockout) with accelerated SAD formation as a tool for genetic manipulation, we clearly demonstrated that age-dependent defect of phagocytosis at maturation stage in astrocytic drives SAD accumulation, synaptic injury and cognitive deficits. Collectively, our findings establish SAD as a mechanistic link between astrocyte senescence and synaptic damage, underscoring the critical role of astrocytic phagocytic function in preserving synaptic homeostasis and cognitive function during aging.

**Significance Statement:** This study reveals that impaired phagocytic maturation in senescent astrocytes leads to formation of SAD and impaired synaptic plasticity, identifying hippocampal astrocyte senescence as a key contributor to age-related cognitive decline.

## Introduction

The progressive decline in synaptic quantity and activity constitutes a core feature of brain aging, which is closely associated with cognitive deterioration. As pivotal regulators of the central nervous system, astrocytes serve as foundational elements in neural network function by maintaining synaptic homeostasis, modulating neurotransmitter balance, and supporting neuronal metabolism^1–3^. Hippocampal astrocytes warrant particular attention—as a structural component of tripartite synapses, which directly participate in the precise regulation of learning- and memory-related neural circuits through dynamic interactions between their processes and pre-/post-synaptic membranes^4^. However, while mechanisms underlying neuronal aging have been extensively studied, significant knowledge gaps remain regarding the functional evolution of astrocytes during aging and their pathological contributions.

Recent studies employing single-cell analysis technologies have revealed regional heterogeneity among astrocytes, suggesting the existence of functionally specialized subtypes in specific brain regions^5^. Nevertheless, these investigations rarely delineate the functional roles of particular astrocyte subtypes within defined brain areas, let alone their specific contributions to health and disease. Astrocytes in hippocampus actively phagocytose synapses to maintain synaptic homeostasis; however, whether specific subtypes of astrocytes are responsible for synaptic clearance in the hippocampus and the alterations in this process during aging or disease conditions remain elusive.

The existence of clustered protein aggregates in the hippocampus has been independently reported by several groups^6–9^. Nevertheless, the nature of these protein aggregates and their impact on hippocampal function remain poorly understood. In this study, we reveal that the clustered protein aggregates in the hippocampus specifically form at aging astrocytic processes (referred to as Senescent Astrocytic Deposits, SAD), and SAD formation is dramatically accelerated in a mouse model (*Nrbf2* knockout mouse) exhibiting dysfunctional phagocytosis. Through genetic approaches, including astrocyte-specific *Nrbf2* knockout and re-introduction, coupled with single-nucleus RNA sequencing (snRNA-seq) and spatial transcriptome (ST) analysis, as well as *in vivo*/*in vitro* phagocytosis assays, we further illustrate that SAD formation is driven by interruption of astrocytic phagocytic function at maturation stage, and contributes to abnormal tripartite synapse structure, loss of synaptic plasticity, and memory damage.

These findings provide novel insights into how astrocyte senescence triggers hippocampal network dysfunction through impairing synapse phagocytosis. This work lays a theoretical foundation for developing glial function-based diagnostic and therapeutic strategies against cognitive impairment.

## Results

### SAD are age-dependent aggregates localized to astrocytic processes

Previous studies have shown that *Nrbf2* knockout (*Nrbf2* KO) mice develop age-related cognitive decline, yet neuronal survival remains intact even at advanced ages^10,11^, suggesting mechanisms beyond direct neuronal regulation contribute to memory deficits. Interestingly, we identified a striking accumulation of clustered p62-positive (p62^+^) protein aggregates in the hippocampus of *Nrbf2* KO mice compared to wild-type (WT) controls (Fig. 1a and Fig. S1a,b), correlating with progressive behavioral impairments characteristic of cognitive impairment. These p62-positive aggregates are located in the hippocampus, accumulate with age, appear earlier and more numerous in Nrbf2 KO mice than in WT mice (Fig. 1b), and can be stained with several antibodies (Table. 1).

**Fig.1.**
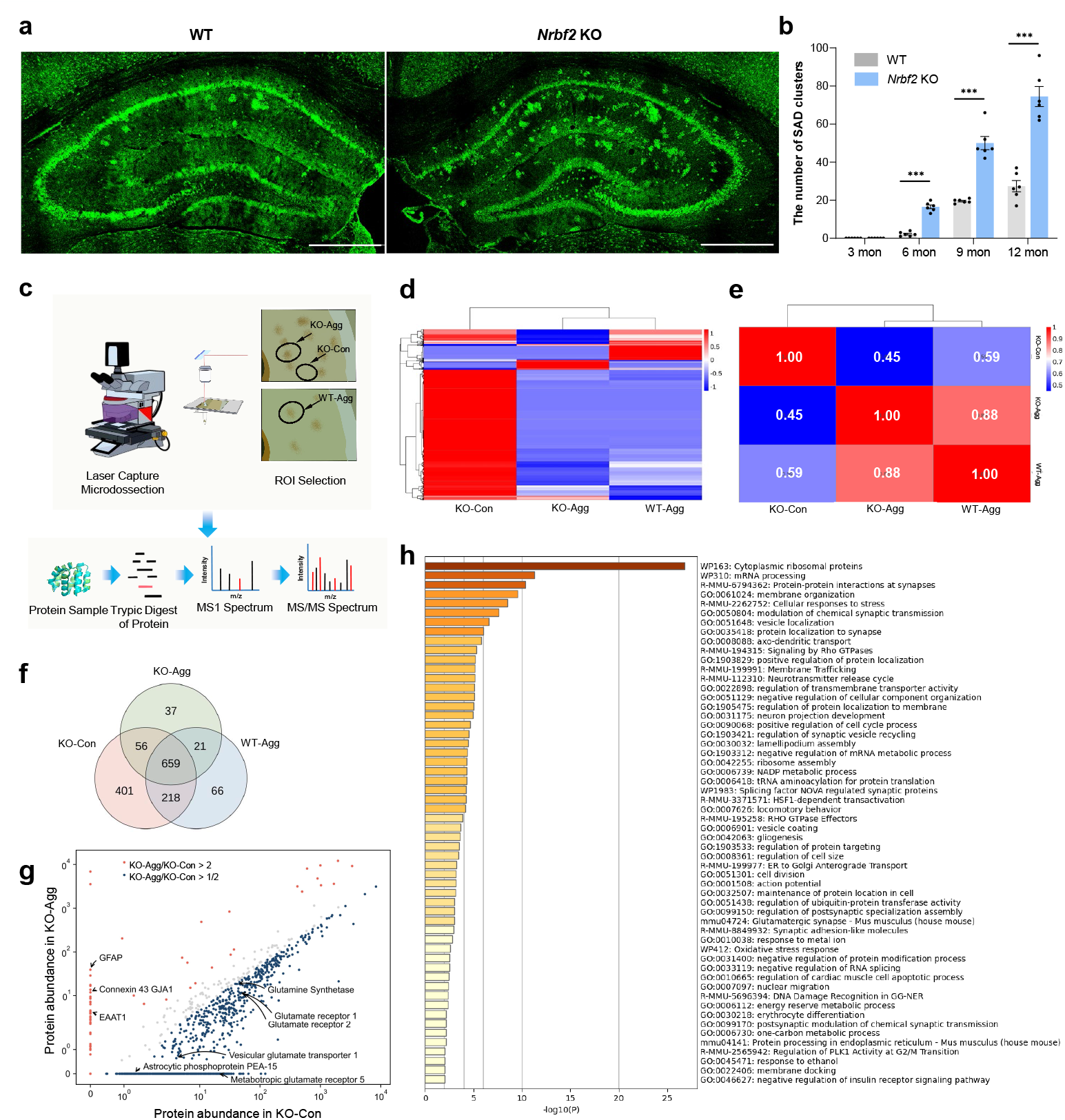
Spatial proteomic analysis reveals that age-dependent p62^+^ aggregates are associated with astrocytes. **a**, Representative confocal images of p62^+^ aggregates (green) in the hippocampus of 12-month-old *Nrbf2* KO and WT mice. Scale bar: 500 μm. **b**, Statistical analysis of SAD numbers in hippocampus of *Nrbf2* KO and WT mice across age groups (n = 6, two-tailed t-test, ^***^p<0.001, data expressed as mean ± SEM). **c**, Schematic workflow of LCM-MS for isolating SAD-containing hippocampal regions. Groups were divided as follows: KO-Agg, hippocampal regions with p62^+^ aggregates from aged *Nrbf2* KO mice; KO-Con, hippocampal regions adjacent to those with p62^+^ aggregates but without p62^+^ aggregates from *Nrbf2* KO mice; WT-Agg, hippocampal regions with p62^+^ aggregates from WT mice. **d**, Comparison of proteomic composition in different groups. **e**, Correlation analysis of proteomic profiles across experimental groups. **f**, Venn diagram of protein overlaps between groups. **g**, Scatter plot of protein expression levels in KO-Agg versus KO-Con groups. **h**, Gene Ontology (GO) cluster analysis of differentially expressed proteins in KO-Agg and KO-Con groups. Data processing: https://metascape.org/gp/index.html.

To define the composition of these p62^+^ aggregates, we performed laser capture microdissection-coupled mass spectrometry (LCM-MS) (Fig. 1c) and revealed highly consistent protein profiles across genotypes in p62^+^ aggregate regions from *Nrbf2* KO and WT mice, whereas marked differences were observed between p62^+^ aggregate and p62^+^ free regions in *Nrbf2* KO mice (Fig. 1d-f) suggesting homogeneity of p62^+^ aggregates across normal aging and *Nrbf2* KO conditions. Differential protein analysis further revealed significant enrichment of astrocytic markers (e.g., GFAP, EAAT1) and synaptic-related proteins in p62^+^ aggregate regions compared to p62^+^ free regions (Fig. 1g and h).

To further characterize the spatial correlation of specific cell types with p62^+^ aggregates, we performed snRNA-seq combined with ST on hippocampal tissues from aged *Nrbf2* KO mice to precisely map gene expression patterns across hippocampal subregional niches (Fig. 2a). The results of gene set enrichment analysis (GSEA) on differentially expressed genes (DEGs) between *Nrbf2* KO mice and WT mice, derived from single-cell sequencing data, demonstrated changes in signaling pathways associated with phagocytosis and neural activity (Fig. S2a,b). The intersection analysis of DEGs between *Nrbf2* KO and WT mice across distinct cell types revealed that Nrbf2 exhibits cell type-specific regulatory functions (Fig. S2d). snRNA-seq of hippocampal astrocytes identified three distinct subclusters (Astro.0, Astro.1, and Astro.2) (Fig. 2b,c). Astro.1 was markedly expanded in *Nrbf2* KO mice, consistently observed in both male and female subjects (Fig. 2c and Fig. S2e). Functional enrichment analysis of its marker genes revealed associations with cell senescence, oxidative stress responses and brain development (Fig. 2d,e). To further infer the functional alterations in Astro.1, we used MEGENA to construct a specific gene co-expression network within this subcluster and identified high expression of *Gfap, Apoe* and *Clu* (Fig. S2f,g). Expression analysis of marker genes in Astro.1 further confirmed the elevated expression of these three genes, with ApoE showing higher expression in *Nrbf2* KO mice, while *Gfap* exhibited no significant difference between WT and *Nrbf2* KO mice (Fig. 2g-i and Fig. S2h-j). These findings suggest that Astro.1 may represent a reactive astrocyte population associated with senescence.

**Fig.2.**
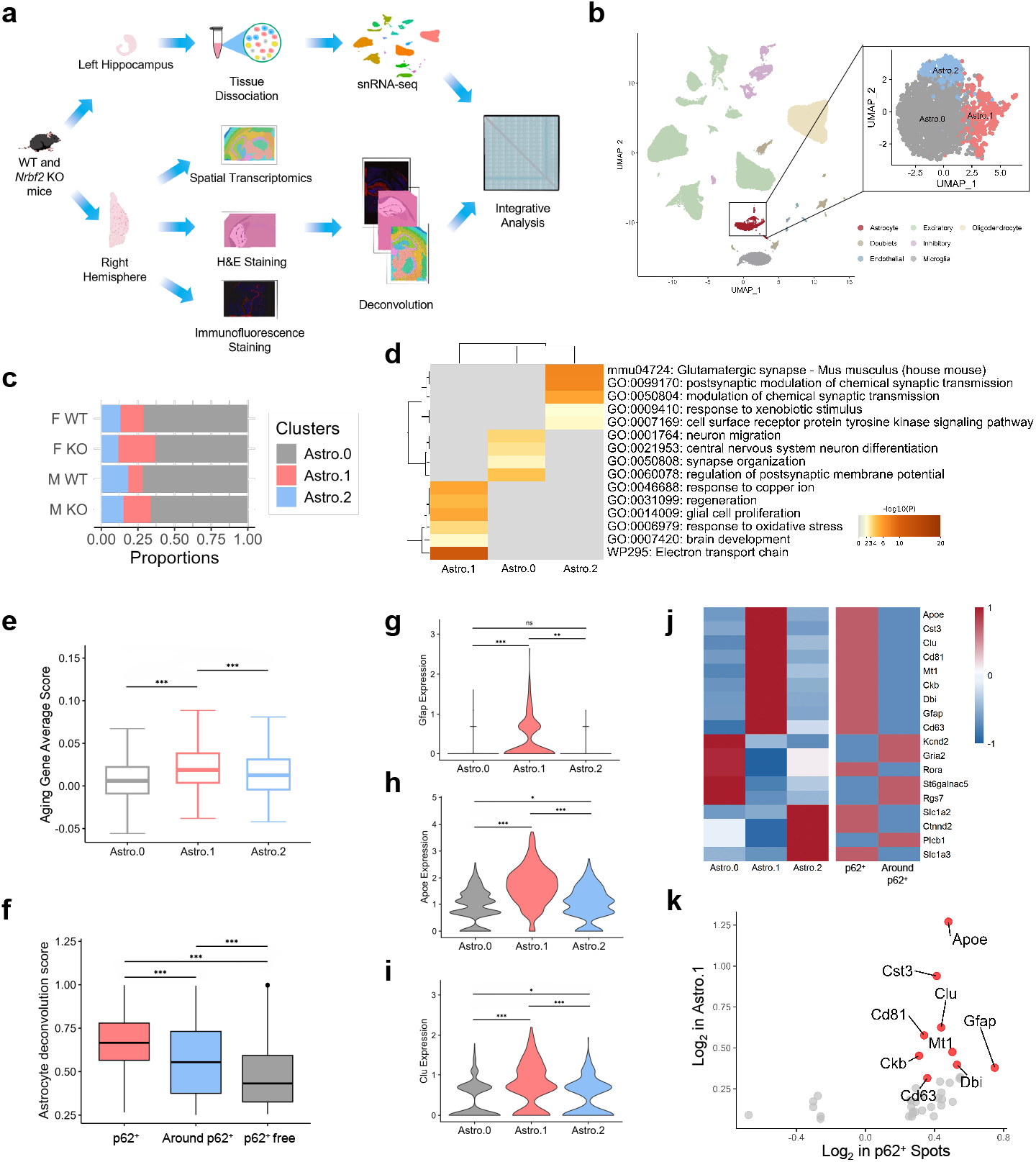
snRNA-seq and spatial transcriptomics analysis reveal that p62^+^ aggregates are associated with a the hippocampus. **a**, Schematic illustration of the snRNA-seq combined with ST experiments. **b**, UMAP visualization of snRNA-seq data. The left panel shows the distribution of major cell types identified in the study, colored as indicated in the legend. The right panel provides a detailed UMAP plot of the astrocyte subcluster, revealing three distinct subclusters (Astro.0, Astro.1, and Astro.2). **c**, Bar plot showing the proportions of each astrocyte subcluster in WT and *Nrbf2* KO hippocampus. **d**, The heat map showing functional pathways enriched by astrocyte subcluster-specific markers. **e**, Boxplots showing age-related gene set scores in different astrocyte subclusters. **f**, The box plots showing the distribution probability of astrocytes in each spot inferred based on the snRNA-seq data. **g-i**, Analysis of the expression level of Astro.1 marker gene. The violin plot shows the expression levels of genes in different astrocyte subclusters. **j**, The heat map showing the relative expression levels of typical marker genes in different astrocyte subclusters and different spot groups. **k**, The scatter plot showing the Astro.1-specific markers that show significant differential expression in p62^+^ aggregate spots. The abscissa represents the expression change fold of these genes in p62^+^ aggregate spots, and the ordinate represents the expression change fold of these genes in Astro.1.

In the analysis of ST data, regions were divided into p62^+^ aggregate regions and p62^+^ free regions by integrating the spatial locations of spots from immunohistochemistry, immunofluorescence (IF), and ST based on p62 fluorescence intensity (Fig. 2a and Fig. S3a,b). The astrocyte deconvolution score demonstrated a high abundance of astrocytes in p62^+^ aggregate regions, which was positively correlated with p62 fluorescence intensity (Fig. 2f). More importantly, in p62^+^ aggregate regions, there was a higher accumulation of Astro.1 compared to Astro.0 and Astro.2 (Fig. S3c-e). Differential expression analysis revealed a significant overlap between genes upregulated in p62^+^ aggregate regions and marker genes of Astro.1, including Gfap and Apoe (Fig. 2j,k). Moreover, the spatial distribution of these marker genes closely matched the pattern of p62 fluorescence intensity (Fig. S3f). These results highlighted the intimate relationship between the p62^+^ aggregates and astrocytes, indicating that p62^+^ aggregate-associated astrocytes are likely a distinct population of activated, senescence-associated astrocytes that are ApoE-high expression.

To validate this astrocyte-specific signature, we performed multiplex IF staining. The results showed that these p62+ aggregates colocalized well with astrocytic processes, and were clustered perfectly within the territory of a single astrocyte (Fig. 3a-c and Video 1). Meanwhile, systematic interrogation of other CNS cell types—including neurons marked by NeuN/β3Tubulin, microglia marked by Iba1, oligodendrocytes marked by CNPase and endothelial cells marked by CD31— showed no spatial correlation with p62+ aggregates (Fig. S4a-d). These results demonstrated that p62^+^ aggregates are uniquely and exclusively linked to astrocytes.

**Fig.3.**
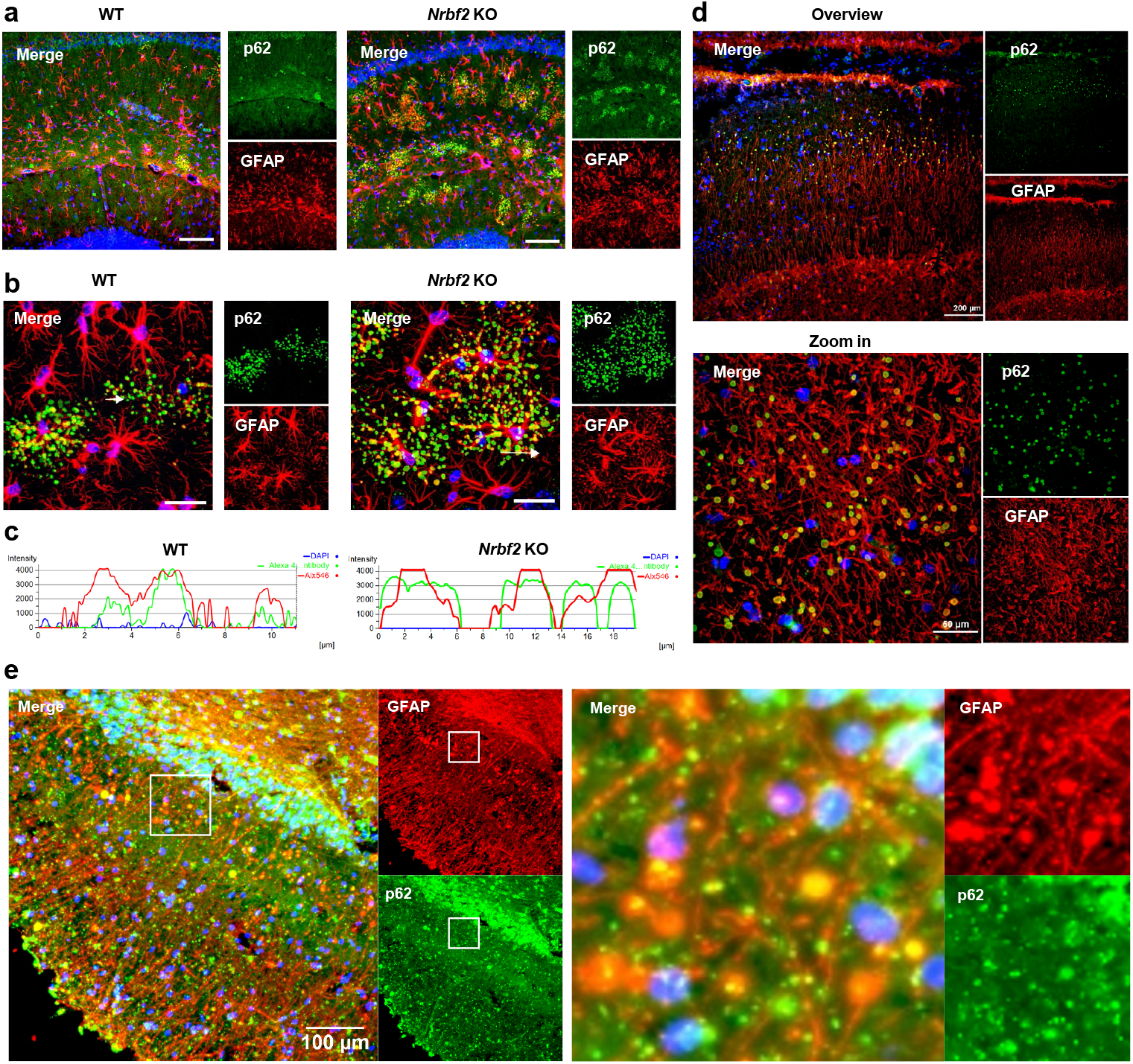
SAD localize at astrocytic processes in hippocampus of aged mice, macaque and human. **a**, Representative confocal images showing co-localization of SAD (p62, green) with astrocytic marker (GFAP, red) in the WT and *Nrbf2* KO mice hippocampal CA1 region. Nuclei stained with DAPI (blue). Scale bar: 100 μm. **b**, Representative 3D-stacked confocal images showing co-localization of SAD (p62, green) with astrocytic processes (GFAP, red) in the WT and *Nrbf2* KO mice hippocampal CA1 region. Nuclei stained with DAPI (blue). Scale bar: 20 μm. **c**, Fluorescence quantitative analysis showing well colocalization of SAD with astrocytic processes. The quantitative position is marked by arrows in panel b. **d**, Representative confocal images of SAD (p62, green) co-localized with astrocytes (GFAP, red) in aged (30-year-old) macaque hippocampus. Nuclei stained with DAPI (blue). **e**, Representative confocal images of SAD (p62, green) co-localized with astrocytes (GFAP, red) in human hippocampus. Nuclei stained with DAPI (blue). Scale bar: 100 μm.

Strikingly, analogous p62+ and GFAP+ astrocytic processes-associated aggregates were observed in the hippocampus of 30-year-old aged macaque, but were absent in younger individuals (Fig. 3d and Fig. S5a). It is worth noting that although these p62+ aggregates in the macaque brain were still located in the CA region of the hippocampus and similarly localized to the terminals of GFAP-positive astrocytes, the morphology of p62+ aggregate-containing astrocytes exhibits a fibrous arrangement. This differs from the astrocytic morphology observed in the mouse brain. These uniquely shaped astrocytes do not colocalize with NeuN-positive or neurofilament-positive cells, thereby excluding the possibility of their neuronal identity (Fig. S5b,c). Most significantly, we observed structurally similar GFAP-positive aggregates in the hippocampal CA region of aged human brains (Fig. 3e). These aggregates formed a near-parallel layer following the trajectory of the pyramidal cell layer, mirroring the spatial organization observed in macaques. This cross-species conservation suggests these structures represent a critical, previously unrecognized component of brain aging processes, which we term Senescent Astrocytic Deposits (SAD).

### SAD disrupt tripartite synapses via ultrastructural disorganization

We next characterized the morphological characteristics of SAD-containing astrocytes. Morphometric analysis demonstrated that SAD-containing astrocytes exhibited significantly larger territorial areas compared to normal astrocytes, likely resulting from SAD formation (Fig. 4a,b). Furthermore, SAD generation led to reduced structural complexity in astrocytes, manifested by decreased branching patterns (Fig. 4c). Transmission electron microscope (TEM) showed that SAD formed clusters and were significantly more abundant in the hippocampus of *Nrbf2* KO mice than in WT controls (Fig.4d and S6a). 3D reconstruction revealed that SAD manifested as irregular, electron-dense aggregates (observed maximum diameter: 3880 nm) (Fig. S6b) localized to astrocytic processes, devoid of membranous encapsulation. Based on their progressive compaction and size, we categorized SAD maturation into three distinct stages: nascent aggregates displayed loosely packed fibrillar material; intermediate-stage SAD developed a condensed core surrounded by fragmented vesicles; and mature SAD exhibited a densely compacted core with pronounced vacuolation in adjacent processes regions (Fig. 4e (ii)-(iv)).

**Fig.4.**
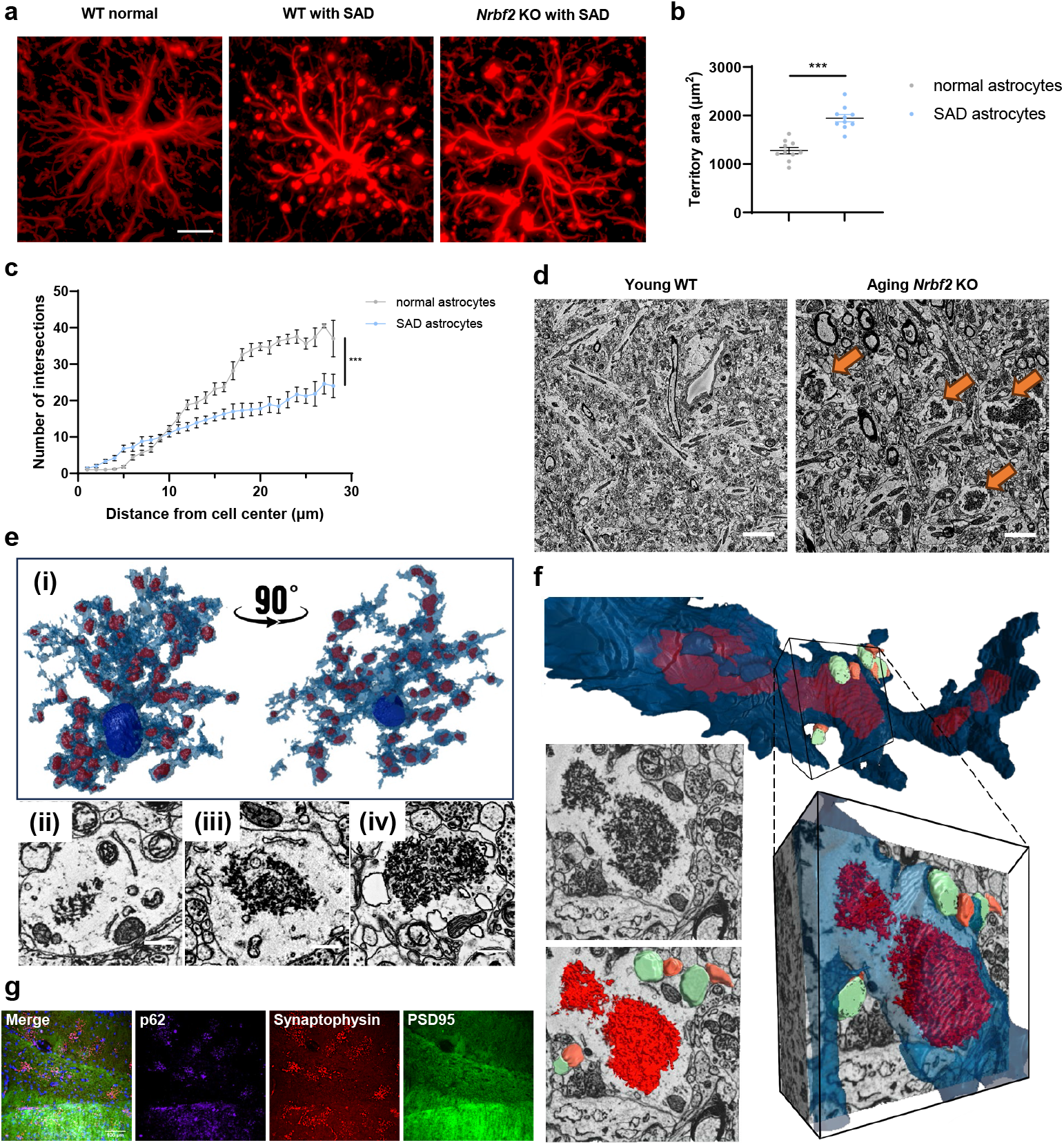
The 3D EM reveals that SAD localize to astrocytic processes and causes abnormal astrocyte morphology. **a**, Representative 3D-stacked confocal images of astrocytes (GFAP, red) without or with SAD from aged WT mice, and astrocytes with SAD from aged *Nrbf2* KO mice. Scale bar: 10 μm. **b**, Scatter plots of territory area in the normal astrocytes and SAD astrocytes (n = 10 cells/group, from 4 mice). **c**, The number of intersections in the normal astrocytes and SAD astrocytes (n = 10 cells/group, from 4 mice). **d**, TEM overviews of SAD distribution (arrows) in young WT versus aged *Nrbf2* KO mice hippocampus. Scale bars: 1 μm. **e, (i)** 3D-EM reconstruction of SAD-containing astrocyte. **(ii)-(iv)** Representative EM images of SAD at three stages. **(ii)** Stage 1: Nascent SAD – Electron-lucent, loosely packed aggregates; astrocytic processes exhibit irregular, amoeboid contours; sparse adjacent synapses with low synaptic vesicle density. **(iii)** Stage 2: Intermediate SAD – Aggregates display a centralized electron-dense core; astrocytic processes adopt rounded morphology; increased synaptic vesicle clusters in neighboring synapses. **(iv)** Stage 3: Mature SAD – Aggregates become hyperdense with a vacuolated central core; astrocytic processes appear spherical and vacuolated. Scale bar: 1 μm. **f**, 3D EM reconstruction of SAD-containing astrocytic processes engulfing synapses. Red: SAD; green: pre-synapse; orange: post-synapse. **g**, Representative confocal images showing the spatial correlation of presynaptic protein (Synaptophysin, red), postsynaptic protein (PSD95, green) with SAD (p62, purple). Nuclei stained with DAPI (blue). Scale bar: 100 μm.

Notably, 3D-EM reconstruction of an entire astrocyte revealed that SAD were specifically localized to astrocytic processes and were not observed outside astrocytes (Fig. 4e (i) and Video 2). These SAD-containing processes were ubiquitously observed and frequently engaged in synaptic engulfment. Surrounding the SAD aggregates, tripartite synaptic structures were frequently observed, comprising the SAD-bearing astrocytic process, presynaptic terminals, and postsynaptic elements (Fig. 4f).Furthermore, we observed that these abnormally enlarged synapses adjacent to SAD-containing astrocytic processes predominantly established contact with astrocytes through their presynaptic rather than postsynaptic components. Immunofluorescence analysis confirmed this structural asymmetry, demonstrating pronounced colocalization of the presynaptic marker synaptophysin with SAD, while the postsynaptic marker PSD95 showed no obvious association (Fig. 4g). These observations lead us to hypothesize that impaired phagosome maturation in SAD-containing astrocytes compromises their synaptic engulfment capacity, resulting in the accumulation and swelling of neighboring synapses.

### Astrocytes govern SAD formation

Next, to further validate the specific role of hippocampal astrocytes in SAD formation, we performed a self-controlled bilateral experiment in *Nrbf2* KO mice with accelerated SAD formation. We injected an adeno-associated virus (AAV) (AAV9::GfaABC1D-*eGfp*-*P2a*-*Nrbf2*-3x*Flag*) promoting NRBF2 overexpression in astrocytes into the CA1 region of the right hippocampus, and a control AAV (AAV9::GfaABC1D-*eGfp*-3x*Flag*) into the same region of the left hippocampus. This setup allowed direct comparison of SAD formation within the same animal, minimizing inter-individual variability (Fig. 5a). Three months after injection, AAV-mediated NRBF2 re-expression in astrocytes significantly reduced SAD burden by 40% compared to the contralateral control (Fig. 5b,c). Strikingly, astrocytes with NRBF2 expression in *Nrbf2* KO mice were nearly devoid of SAD, whereas adjacent uninfected astrocytes retained abundant aggregates as highlighted in yellow circle (Fig. 5d,e), confirming that NRBF2 restoration specifically in astrocytes rescues SAD pathology. Neuronal-specific NRBF2 overexpression by AAV9::Syn-*eGfp*-*P2a*-*Nrbf2*-3x*Flag* in *Nrbf2* KO mice’s hippocampus failed to suppress SAD accumulation (Fig. S7a–d). These results demonstrate that the massive SAD burden observed in *Nrbf2* KO mice is attributable to astrocyte-specific deletion of *Nrbf2*, independent of neuronal involvement.

**Fig.5.**
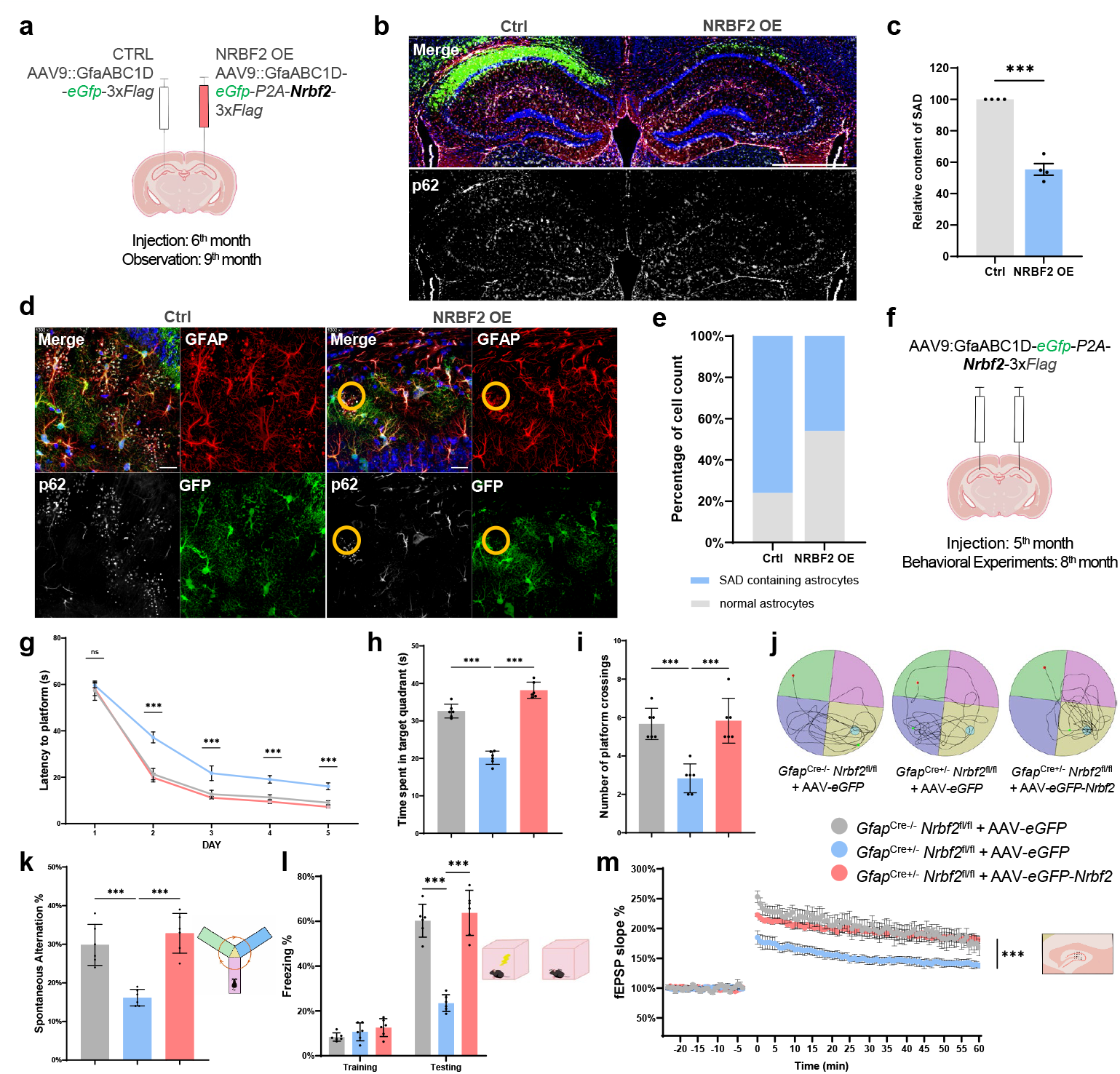
Astrocytic NRBF2 prevents SAD formation, maintains cognitive function and synaptic plasticity. **a**, Schematic of astrocyte-specific NRBF2 rescue experiment. **b**, Representative confocal images of SAD burden in the hippocampus of *Nrbf2* KO mice after injection of NRBF2-overexpressing AAV (AAV9::GfaABC1D-*eGfp*-*P2a*-*Nrbf2*-3x*Flag*) and control AAV (AAV9::GfaABC1D-*eGfp*-3x*Flag*). SAD (p62, white); GFP (green); astrocytes (GFAP, red); nuclei stained with DAPI (blue). Scale bars: 500 μm. **c**, Quantification of SAD area in the bilateral hippocampus after AAV injection (n = 4 mice; two-tailed t-test, ^***^P < 0.0001). **d**, Representative images of SAD expression in astrocytic processes of the bilateral hippocampal CA1 region after injection of NRBF2-overexpressing AAV and control AAV. GFAP (red), GFP-NRBF2/GFP (green), SAD (p62, white), and DAPI (nuclei, blue). Scale bar, 20 μm. **e**, Quantification of the number of SAD-containing astrocytic processes in the bilateral hippocampus after AAV expression (n=50 per group from 4 mice). **f**, Experimental schedules of AAV (*NRBF2* OE) reporter injection and analysis. **g**, The escape latency to platform of each group in the training phase of Morris water maze test (n = 6). **h and i**, Time spent in the target quadrant and the number of platform crossings by each group in the probe trial of Morris water maze test (n = 6). **j**, Representative images of the swimming tracks of each group in the probe trial of the Morris water maze test (n = 6). **k**, The percentage of spontaneous alternations of each group in the Y maze test (n = 6). **l**, Freezing behavior of each group in the contextual fear conditioning test (n = 6). **m**, Electrophysiological results showing that the LTP decline caused by *Nrbf2* deficiency can be rescued in *Gfap*^Cre+/-^ *Nrbf2*^fl/fl^ mice after injection AAV (NRBF2 OE) (n=6 slices from 3 mice). Significance was determined using one-way ANOVA followed by Dunnett’s multiple comparison test in h-i. ^*^P < 0.05, ^**^P < 0.01 and ^***^P < 0.005. Mean ± S.E.M.

To further dissect this astrocyte-specific mechanism, we generated loxP-floxed *Nrbf2* (Nrbf2^*fl/fl*^) mice enabling Cre-dependent conditional knockout. By crossing with Gfap^Cre+/-^ and Aldh1l1^CreERT2+/-^ mice, we established two astrocyte-specific *Nrbf2* knockout mice (Fig. S8a,b). Both *Gfap*^*Cre+/-*^ *Nrbf2*^*fl/fl*^ and *Aldh1l1*^*CreERT2+/-*^ *Nrbf2*^*fl/fl*^ mice exhibited normal astrocyte numbers at 6 months (Fig. S8c,d). However, SAD density in the hippocampus increased dramatically in both *Gfap*^*Cre+/-*^ *Nrbf2*^*fl/fl*^ and *Aldh1l1*^*CreERT2+/-*^ *Nrbf2*^*fl/fl*^ mice compared to littermate controls, which recapitulated that observed in global *Nrbf2* KO mice (Fig. S8e), indicating that functional impairment of astrocyte, due to *Nrbf2* deletion, is sufficient to drive SAD pathology.

### Astrocytic NRBF2 regulates synaptic plasticity and cognitive function

Synaptic plasticity—the activity-dependent modulation of synaptic strength—is a fundamental cellular mechanism underlying learning and memory. Deficits in synaptic plasticity and memory decline are hallmark features of neurodegenerative dementia^12^. While global *Nrbf2* deletion has been shown to impair memory by disrupting synaptic plasticity in mice, the specific contribution of astrocytes to these processes remained undefined^11^. To address this, we subjected comprehensive behavioral and electrophysiological analyses. *Aldh1l1*^CreERT2+/-^ *Nrbf2*^fl/fl^ mice developed observable behavioral deficits by 6 months of age compared to WT littermates (Fig. S8f-i), demonstrating that astrocyte specific *Nrbf2* deletion alone was sufficient to induce behavioral abnormalities in mice, mirroring phenotypes observed in global knockouts. In the Morris water maze (MWM), *Gfap*^Cre+/-^ *Nrbf2*^fl/fl^ mice exhibited prolonged escape latency, reduced target quadrant occupancy, and fewer platform crossings (Fig. 5g-j). They exhibited poor performance in the spontaneous alternation test (Fig. 5k), indicating impaired spatial working memory. In the contextual fear conditioning (CFC) test they showed diminished freezing duration (Fig. 5l), reflecting compromised memory consolidation. AAV9::GfaABC1D-mediated NRBF2 expression selectively in hippocampal astrocytes of 6-month-old *Gfap*^Cre+/-^ *Nrbf2*^fl/fl^ mice for 3 months fully rescued memory impairments (Fig. 5f-l). Electrophysiological recordings from hippocampal slices revealed that *Gfap*^Cre+/-^ *Nrbf2*^fl/fl^ mice exhibited significantly reduced long-term potentiation (LTP), a canonical correlate of synaptic plasticity. Critically, astrocytic NRBF2 reconstitution rescued the fall of LTP in *Gfap*^Cre+/-^ *Nrbf2*^fl/fl^ mice (Fig. 5m), indicating that the loss of synaptic plasticity in this mouse model is solely caused by astrocyte dysfunction. These results establish that astrocytic NRBF2 is indispensable for maintaining synaptic plasticity and learning-dependent behaviors.

### Phagocytic dysfunction of astrocytes drives SAD formation

3D-EM results revealed that astrocytic processes containing SAD frequently exhibited ongoing synaptic engulfment, with phagosomes containing undegraded synaptic debris trapped near SAD margins (Fig. 4f). Strikingly, quantitative analysis revealed profound disruptions in tripartite synaptic architecture near SAD-containing processes, characterized by the volume of synapses adjacent to SAD-containing processes being markedly enlarged compared to that of synapses distal to SAD (Fig. 6a,b and Fig. S6c). This hypertrophy was accompanied by an abnormal accumulation of multiple synaptic structures surrounding individual processes, a phenomenon rarely observed near healthy tripartite synapses (Fig. 6c). These structural anomalies prompted our investigation into impaired phagocytic processes in SAD-containing astrocytes within the hippocampi of both aging and *Nrbf2* KO mice.

**Fig.6.**
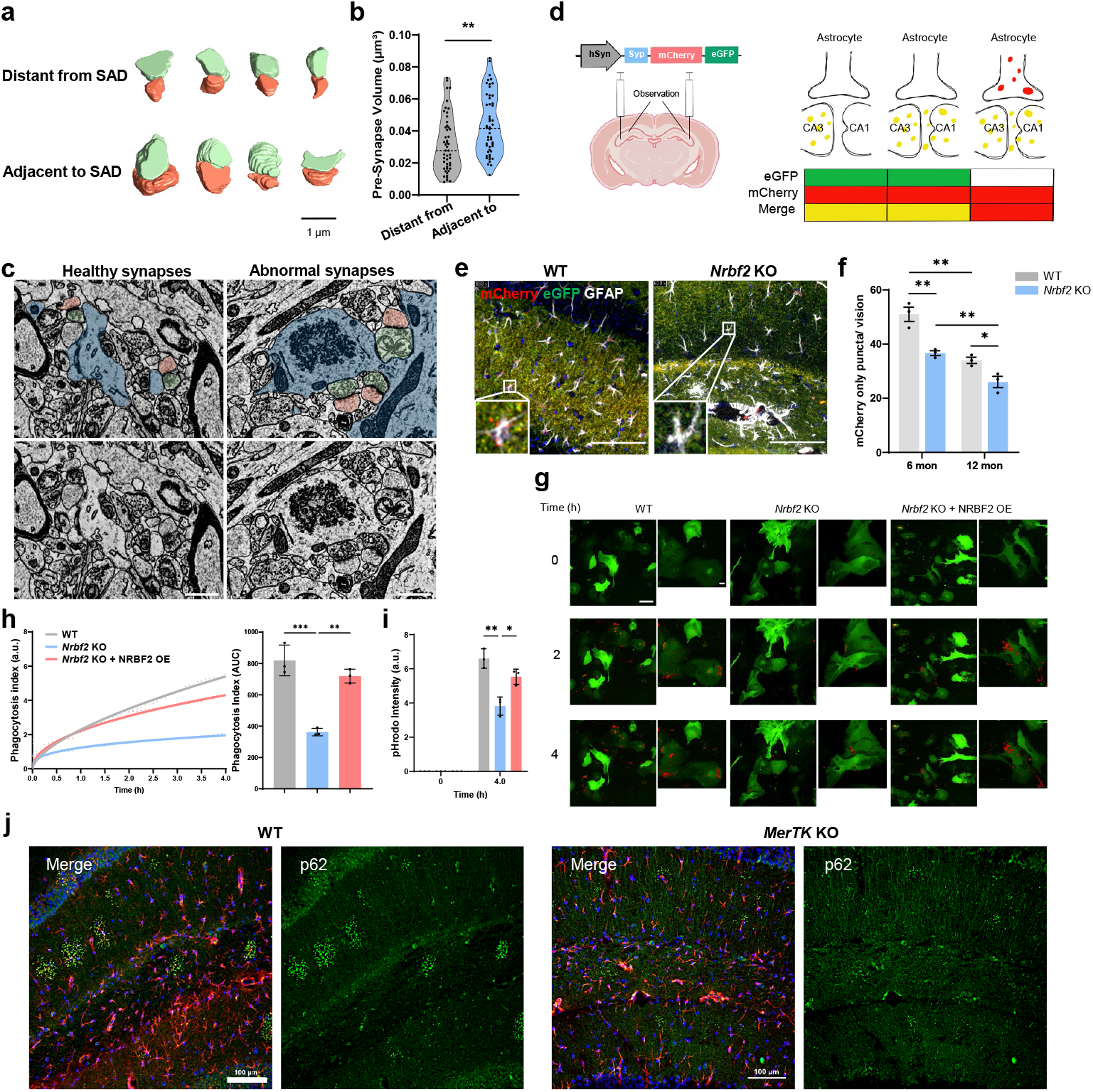
Astrocytic phagocytosis dysfunction specifically at maturation stage drives SAD formation. **a**, Representative 3D-EM reconstruction showing synaptic morphological distant from and adjacent to SAD-containing astrocytic processes. Scale bar: 1 μm. **b**, Quantification of synaptic volumes distant from and adjacent to SAD-containing astrocytic processes (n = 50 per group; two-tailed t-test, ^**^p<0.01). **c**, Representative EM images showing the accumulation of multiple morphologically abnormal synapses around astrocytic processes containing SAD. Scale bar: 1 μm. **d**, Schematic of hSyn-*Syp-mCherry-eGfp* reporter injection. **e**, Representative confocal images of mCherry-only puncta reduction in CA1 astrocytes (mCherry, red; EGFP, green; astrocytes stained with GFAP, white; Nuclei stained with DAPI, blue. Scale bar: 100 μm). **f**, quantification of the mCherry only puncta in different groups. One-way ANOVA followed by Dunnett’s multiple comparison test, ^*^P < 0.05 and ^**^P < 0.01. **g**, Representative images of synaptic engulfment by primary astrocytes drive from WT and *Nrbf2* KO mice overexpressing NRBF2 (primary astrocyte EGFP, green; synaptosome stained with pHrodo Red, red. Scale bar: 50μm, zoom in 10μm). **h**, Quantification of phagocytosis index as arbitrary units for the Time lapse images over 4h, 5min intervals, phagocytosis index is defined as the number of pHrodo red synaptosomes ingested by one primary astrocyte. Quantification of phagocytosis index is shown as area under curve (AUC). **i**, Bar graphs show pHrodo fluorescence intensity expressed as arbitrary units. Ordinary one-way ANOVA in (e) and two-way ANOVA in (f); n = 3 independent experiments; ^**^P<0.01 and ^***^P<0.001. **j**, Representative confocal images of SAD (p62, green) in the hippocampus of 14-month-old WT and MerTK KO mice. GFAP (red), and DAPI (nuclei, blue), Scale bar: 100 μm.

Astrocyte-mediated synaptic elimination in the adult hippocampus is critical for maintaining synaptic connectivity and plasticity^13,14^. Our previous studies demonstrated that NRBF2 is required for phagocytosis both *in vitro* and *in vivo*^15^. We thus investigated whether NRBF2 sustains synaptic plasticity and homeostatic regulation by modulating astrocytic synaptic elimination activity. To assess phagocytic activity, we injected an established dual-fluorescence reporter system hSyn-*Synaptophysin*-*mCherry*-*eGfp* into the hippocampal CA3 region as previously described^13^. Acidification-dependent eGFP quench in phagolysosomes of CA1 astrocytes allowed quantification of mCherry-only puncta, reflecting phagolysosomal degradation (Fig. 6d). In *Nrbf2* KO mice, we observed a significant age-dependent reduction in mCherry-only puncta within the CA1 region compared to that in WT mice (Fig. 6e,f), indicating impaired synaptic phagocytosis. Primary astrocytes isolated from *Nrbf2* KO mice exhibited similar deficits, with reduced phagocytosis of pHrodo-labeled synaptosomes and defective acidification of late endosomes—a critical step for phagosome maturation (Fig. 6g-i; Fig. S9a-c; Video 3). Notably, lysosomal marker LAMP1 and cathepsin B/D expression and baseline acidity remained unaltered (Fig. S9d,e), localizing the defect to phagocytic maturation rather than lysosomal dysfunction.

Our previous work has revealed the role of NRBF2 as an effector of RAB7 to promote phagosome-lysosome fusion^16^. To determine whether the effect of astrocytic NRBF2 on phagocytosis is mediated by regulating RAB7 activity, we assessed the activation of RAB7 in primary astrocytes using the GST-R7BD and Raichu-RAB7 FRET sensor as described previously^16^. The results showed that NRBF2 deficiency significantly reduced the GTP form of RAB7 in primary astrocytes without affecting the total RAB7 expression (Fig. 7c, Fig. S 9f,g). To establish the causative link between phagocytosis impairment and SAD formation, we overexpressed RAB7 in NRBF2-deficient astrocytes and observed effectively restored phagocytic activity and phagosomal acidification (Fig. 7a,b). Similarly, RAB7 overexpression in *Nrbf2* KO hippocampus partially reduced SAD burden as expected (Fig. 7d-f). These findings indicate that astrocytic NRBF2 promotes phagosome maturation by regulating RAB7 activity. In summary, dysfunctional phagocytosis maturation of astrocytes in the *Nrbf2* KO mice acts as a central driver of SAD aggregation.

**Fig.7.**
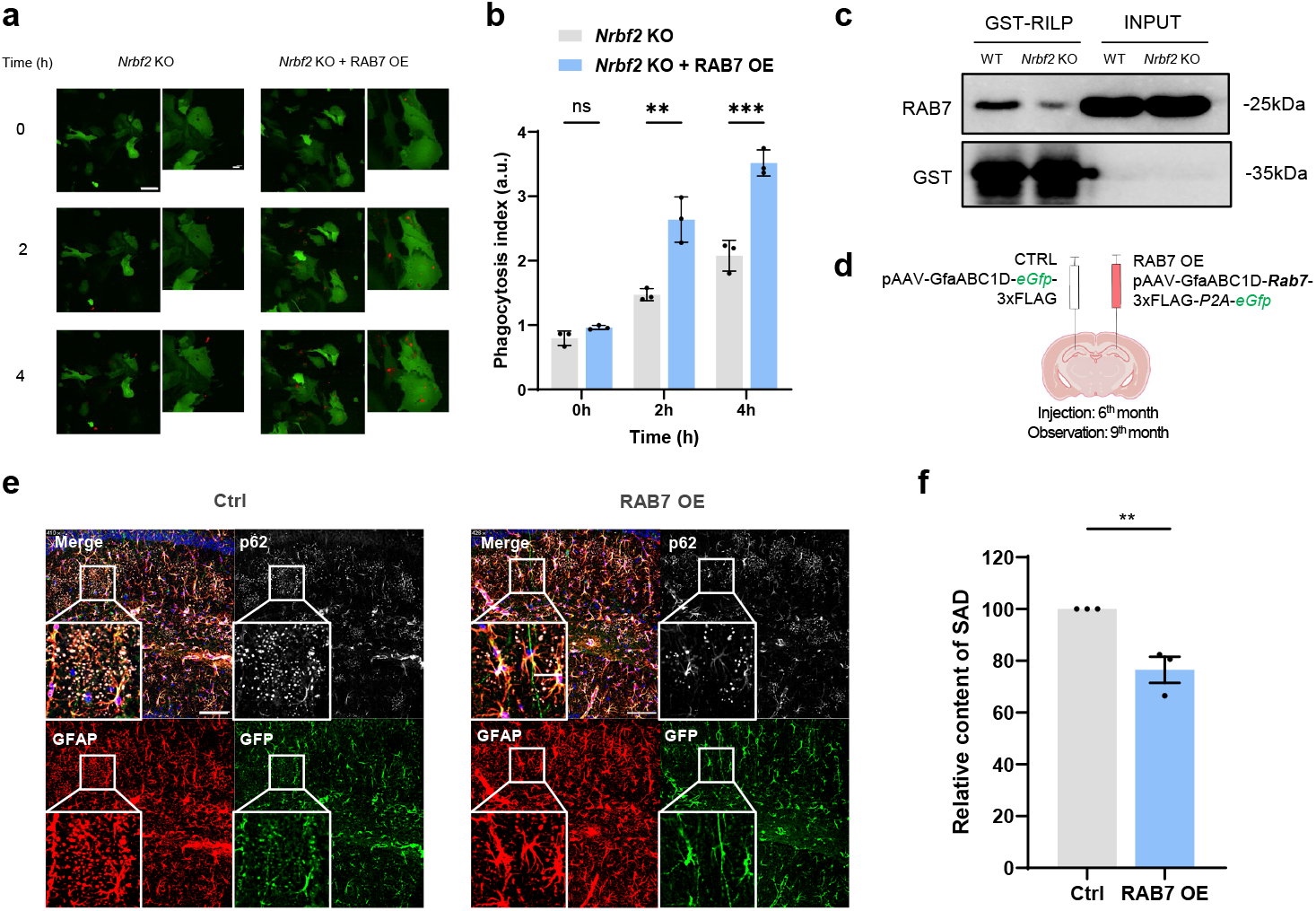
Restoration of impaired phagocytosis alleviates SAD formation. **a**, Representative images of synaptic engulfment by primary astrocytes derived from *Nrbf2* KO mice overexpressing RAB7 and NRBF2 (primary astrocyte EGFP, green; synaptosome stained with pHrodo Red, red. Scale bar: 50μm, zoom in 10μm). **b**, Bar graphs show phagocytosis index expressed as arbitrary units. Ordinary one-way ANOVA in (e) and two-way ANOVA in (f); n = 3 independent experiments; ^**^P<0.01 and ^***^P<0.001. **c**, GST-R7BD pull-down assay indicates that active RAB7 is reduced in *Nrbf2* KO primary astrocyte. The “GST-RILP” indicated pull-down with GST-RILP beads. **d**, Experimental schedules of AAV9-mediaetd *Rab7* OE and analysis. **e**, Representative images of SAD formation in the CA1 region of the hippocampus of *Nrbf2* KO mice after injection of AAV (RAB7 *OE*) and control AAV. Scale bars: 100 μm; **f**, Quantification of SAD area in the bilateral hippocampus after AAV injection (n = 3 mice; two-tailed t-test, ^**^P<0.01).

Phagocytosis can be simply divided into 3 steps: recognition of cargos, internalization of cargos to form phagosome, phagosome maturation and degradation. To further define the role of impaired phagocytosis maturation in SAD formation, we analyzed *MerTK* KO mice, a model in which astrocytic recognition and engulfment of cargos are fundamentally abolished^14^. As expected, we observed a near absence of SAD in the hippocampus of *MerTK* KO mice, which was significantly reduced compared to age-matched WT controls (Fig. 6j). These results demonstrate that SAD generation relies on normal engulfment of synaptic materials, and the impairment of phagosomal maturation in astrocytes.

## Discussion

Brain aging is intricately linked to synaptic dysfunction and cognitive decline, yet the role of astrocyte senescence in these processes has remained enigmatic. Our identification of SAD—hippocampal astrocytic processes localized protein aggregates—provides a mechanistic bridge between astrocyte aging and tripartite synaptic dysfunction. Through cross-species analyses in mice and primates, we demonstrate that SAD accumulation correlates with synaptic loss, ultrastructural disorganization of tripartite synapses, and memory deficits. Crucially, SAD pathology is driven by decreased astrocytic phagocytic activity, while NRBF2 deficiency accelerates this process by disrupting RAB7-dependent phagosome maturation. These findings establish SAD as a conserved biomarker of astrocyte aging and memory decline, and highlight NRBF2-dependent phagocytosis as a critical pathway for maintaining synaptic homeostasis (Fig. 8).

**Fig.8.**
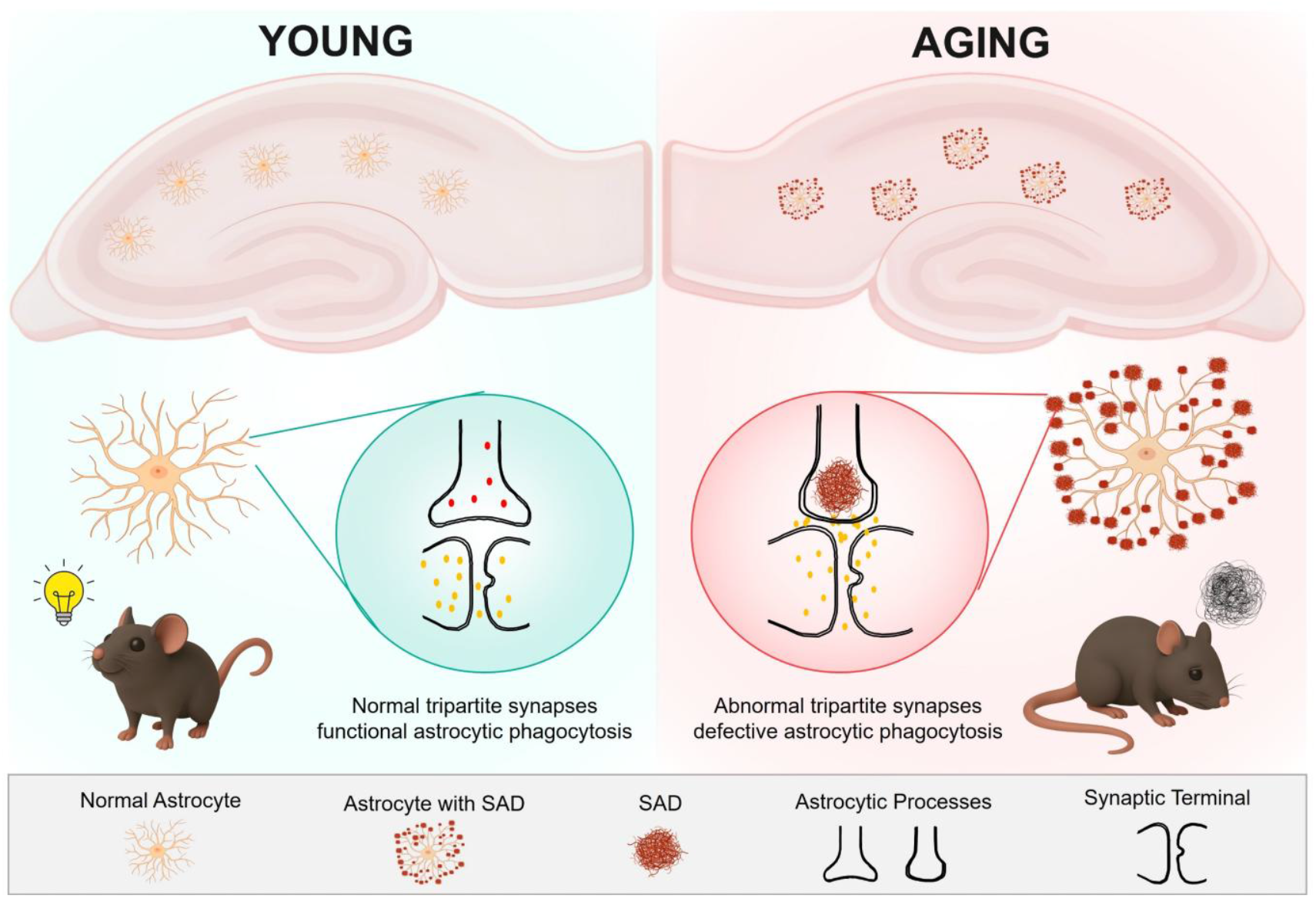
Summary diagram.

Recent advances in astrocyte biology have revealed striking regional heterogeneity^17^, while functional specialization within hippocampal astrocytes and their contributions to aging remain undefined. Unlike the previous view that astrocytes in the brain are largely uniform, astrocytes in different structures and even substructures exhibit genetic and functional diversity. Some studies have also shown the functional specificity of certain astrocytes in the hippocampal region^18–20^. Nevertheless, the specificity of astrocytes in the hippocampus and its associated neural circuits, as well as their contributions to pathological processes, have largely remained unexplored. Our spatial transcriptomic and snRNA-seq analyses identified a distinct astrocyte subpopulation, characterized by upregulation of *Gfap* and *Apoe*. This subpopulation is precisely distributed in the hippocampus and exhibits a unique propensity for SAD formation, linking local phagocytic dysfunction to synaptic pathology. The 3D-EM evidence further demonstrates that SAD physically disrupt tripartite synapses, impairing synaptic morphology and plasticity. Importantly, the conservation of SAD in aged primates and human underscores their translational relevance as a cross-species hallmark of brain aging. Crucially, the data from cross-species samples consistently reveal that SAD exhibit a striking spatial constraint in the hippocampal CA region, the pivotal region for memory, which links SAD accumulation to age-related cognitive decline. These findings not only confirm the functional differentiation of astrocyte subtypes, but also suggest that pathological changes in specific brain regions may originate from the dysfunction of local glial cells.

The tripartite synapse, composed of astrocyte processes, pre- and post-synapse, is fundamental for hippocampal synaptic plasticity and memory. Though past studies mainly focused on the information exchange between astrocytes and synapses, our results indicates that astrocytes actively phagocytose synapse to maintain quality of tripartite synapse. Age-dependent decline in astrocytic phagocytosis function caused SAD accumulation to disrupt tripartite synapse, leading to synaptic terminal hypertrophy and loss of synaptic plasticity. The accumulation of tripartite synapses surrounding SAD-containing astrocytic processes of the ApoE-high expression astrocyte subtype highlights the implication of this astrocyte subtype in the tripartite synapse dysfunction and memory defect during aging. Further analysis of this astrocyte subtype would expand our understanding of how regional glial heterogeneity governs tripartite synaptic damage during aging and why hippocampal circuits are preferentially targeted.

The clearance capacity of astrocytes is essential for maintaining brain homeostasis^21^. The formation of SAD is dependent on normal astrocytic engulfment, followed by impaired degradative activity. SAD contain a range of non-astrocytic proteins, including synaptic proteins and matrix proteins, suggesting a direct association with the phagocytosis process. By over-expressing RAB7, a key protein in phagosome maturation, we observed a reduction in SAD formation, thereby confirming that deficiencies in the endolysosomal system drive the formation of SAD. These findings align with human studies showing age-dependent accumulation of synaptic proteins in astrocytes^22^, emphasizing phagocytic clearance—rather than engulfment—as the critical bottleneck in aging.

Our previous work has established the role of NRBF2 in autophagy regulation^23^, raising the concerns regarding the involvement of autophagy deficits in SAD formation. Notably, accumulation of SAD was not obvious in several mice stain with canonical autophagy genes deletion (*Atg7, Ulk1/2, Fip200*) in astrocytes^24,25^, implying that dysfunctional autophagy may not directly contribute to SAD formation.

Furthermore, the role of NRBF2 in astrocytes seems to be particularly crucial. Astrocyte-specific NRBF2 reconstitution not only reduced SAD burden but also restored synaptic plasticity and memory performance, demonstrating the reversibility of age-related cognitive deficits. Notably, this effect was absent with neuron-specific NRBF2 overexpression, suggesting an astrocyte-dominant role in regulating synaptic plasticity and memory during aging. Given aging represents the primary risk factor for neurodegenerative disorders, future studies should explore whether SAD contributes to neurodegenerative pathologies such as Alzheimer’s disease beyond general aging processes.

We propose that impaired phagocytic degradation in aging astrocytes leads to the pathological retention of synaptic debris and other proteins within processes to form SAD which results in accumulation of dysfunctional synapses and impaired cognitive function. In summary, our work delineates a hippocampus-specific astrocyte subpopulation whose phagocytic decline drives SAD accumulation and cognitive impairment during aging. By identifying NRBF2 as an essential regulator of astrocytic phagocytosis and SAD formation, we provide a mechanistic framework for targeting glial dysfunction in age-related cognitive decline. These insights open avenues for diagnostics and therapies aimed at preserving synaptic integrity and cognitive resilience in the aging brain by modulating astrocytic phagocytosis activity.

## Methods and Reagents

### Mice

All animal experiments were approved by the University of Macau Ethics Committee and complied with relevant ethical guidelines (approval number: UMARE-28-2024). Animals were group-housed under a 12-hour light/dark cycle with ad libitum access to food and water. The ambient temperature was maintained at 24°C with a relative humidity of 60% ± 5%. For experiments involving mutant mice, littermates were used as controls. All mice were randomly assigned to experimental groups without sex-based distinctions.

#### Generation of *Gfap*^*Cre*+/-^ *Nrbf2*^*fl/fl*^ and *Aldh1l1*^*Cre*+/-^ *Nrbf2*^*fl/fl*^ mice

*Gfap*^*Cre+/-*^ *Nrbf2* ^*fl/fl*^ and *Aldh1l1*^*Cre+/-*^ *Nrbf2*^*fl/fl*^ mice were generated by crossing homozygous *Nrbf2*-floxed mice (*Nrbf2*^*fl/fl*^, C57BL/6 background) with either *Gfap*^*Cre*+/-^ or *Aldh1l1*^*CreERT2* +/-^ driver lines. Genotypes were confirmed by PCR screening of tail DNA using allele-specific primers: Nrbf2: ①(5’arm) ATTTGTCACGTCCTGCACGA

TAGCTTCTTGGTTAGGTGTGAGAC

②(3’arm) TTCCCAAGACCCTCATGGTAG

CACAGGGTTCAATTTCCAGTACC.

Gfap-iCre: ①5’arm GGGCAGTCTGGTACTTCCAAGCT

TCAGCAACGCTGGAGAATCCC

②WT CAGCAAAACCTGGCTGTGGATC

ATGAGCCACCATGTGGGTGTC

Aldh1l1-CreERT2: ①5’arm GCCCTGTATGTCAGTGACAAACTG

CTGGTTCTTTCCGCCTCAGA

②WT GCCCTGTATGTCAGTGACAAACTG

CAGGGTACAGGAGAAGTATCACAGTC

Recombination efficiency and spatial specificity of NRBF2 depletion in astrocytes were validated by extracting primary hippocampal astrocytes and performing WB analysis of GFAP. All experiments used littermate controls, and mice were randomly assigned to groups without sex-based distinctions.

### Immunofluorescence and Immunohistochemistry

Mouse brain samples were fixed in 4% paraformaldehyde (Wako Chemicals), followed by graded sucrose dehydration. Tissues were embedded in OCT compound and sectioned at 20 μm. Free-floating sections were blocked with goat serum and permeabilized with 0.3% Triton X-100 at room temperature. Sections were then incubated overnight at 4 °C with primary antibodies at appropriate dilutions. After washing with PBST (PBS containing 0.1% Tween 20), sections were incubated at room temperature with fluorophore-conjugated secondary antibodies or enzyme substrates for immunohistochemistry. Images were acquired using a Nikon A1R confocal microscope.

Brain tissues of 13-year-old and 30-year-old rhesus macaques were retrieved from National Resource Center for Non-Human Primates, Kunming Institute of Zoology (https://nhp.kiz.ac.cn/), which offers regular tissues of non-human primates. All samples were collected following a protocol approved by the Institutional Animal Care and Use Committee (IACUC) of Kunming Institute of Zoology (IACUC-PE-2024-12-003).

### laser capture microdissection-coupled mass spectrometry (LCM-MS)

Sample Preparation: Formalin-fixed paraffin-embedded (FFPE) tissue sections were deparaffinized with xylene, dehydrated with absolute ethanol, and vacuum-dried. Dried tissue was homogenized in lysis buffer (50 mM ABC, pH 8.0, 75 mM NaCl, 0.1 M DTT) and incubated at 99°C for 60 min. After cooling, sodium deoxycholate (DOC) was added to 2% final concentration. Samples were sonicated (pulsed, 99% power, on ice), centrifuged (16,000 × g, 10 min), and the supernatant was subjected to filter-aided sample preparation (FASP) using 30 kDa molecular weight cut-off filters. Proteins were washed with 50 mM ABC buffer, digested on-filter with trypsin (1:100 enzyme-to-protein ratio) in 50 mM ABC containing 0.5% DOC at 37°C for 4–18 hours. Peptides were eluted with ABC buffer and acidified with formic acid prior to desalting.

LC-MS/MS Analysis: Peptides were separated on a reversed-phase C18 column (150 µm × 15 cm, 1.9 µm particles, 120 Å pore) using an Easy-nLC 1200 system with a gradient of 10–95% mobile phase B (80% acetonitrile, 0.1% formic acid) over 60 min at 600 nL/min. Eluted peptides were ionized via nano-electrospray and analyzed on an Orbitrap Fusion Lumos mass spectrometer operated in data-dependent acquisition (DDA) mode. Full MS scans (m/z 350-1550) were acquired in the Orbitrap at 120,000 resolution. The most intense ions (top speed mode) were selected for HCD fragmentation (32% NCE) and MS/MS spectra were acquired in the Orbitrap at 7,500 resolution.

Data Analysis: Raw files were searched against the UniProt Mouse reviewed database using Proteome Discoverer (v2.2) and SEQUEST HT with Percolator validation. Search parameters included: 20 ppm precursor tolerance, 0.05 Da fragment tolerance, trypsin specificity (⩽2 missed cleavages), carbamidomethylation (C) as fixed modification, and variable modifications for oxidation (M), N-terminal acetylation, and lysine acetylation. Peptide spectral matches and protein identifications were filtered to a 1% false discovery rate (FDR).

### Transmission electron microscope (TEM) and 3D-EM reconstruction

Sample Preparation: Mice were transcardially perfused with ice-cold fixative (2.5% glutaraldehyde [EMS, 16000], 4% paraformaldehyde [EMS, 15700] in 0.1 M phosphate buffer, pH 7.4). Brains were dissected, sectioned at 100 µm using a vibratome (Leica, VT1200 S), and hippocampal slices were post-fixed in 2.5% glutaraldehyde at 4°C overnight. After PB rinses, slices underwent sequential treatments: 2% OsO_4_ (Serva)/3% potassium ferrocyanide (Sigma-Aldrich, 455989) at 4°C, 1% thiocarbohydrazide (Sigma-Aldrich, 223220) at RT, 2% OsO_4_ at RT, and 2% uranyl acetate at 4°C overnight. Samples were then incubated in lead aspartate solution (lead nitrate [Sigma-Aldrich, 203580], pH 5.4), dehydrated through graded ethanol/acetone, and embedded in resin (SPI, Epon812) with gradient infiltration. Polymerization occurred at staged temperatures (37°C→45°C→60°C).

Data Collection: Trimmed samples (quadrangular frustum pyramids) were coated with 40nm Platinum using a Vacuum Evaporator (Leica, EM ACE600) and imaged on an ultramicrotome (Zeiss, Volutome) via Serial Block Face. Parameters: 60×60 µm field, 35 µm z-stack, 7 nm pixels, 70 nm slices, 1.5 µs dwell time, 2× averaging.

Volume Reconstruction: Image stacks were reconstructed in Amira (Thermo Fisher) and Dragonfly (Comet Technologies Canada). Astrocytes containing SABs were identified using a deep learning model (Amira) trained on manual annotations (criteria: SAB presence, clear cytoplasm), with post-hoc correction. Synapses were detected in Dragonfly using a pre-trained model (criteria: presynaptic vesicles, synaptic cleft, postsynaptic density). The Allen Brain Atlas hippocampus mesh was aligned to the EM dataset location for 3D visualization.

### AAV stereotaxic surgery

Mice were anesthetized with isoflurane (5% induction in a sealed chamber, 1.5–2% maintenance via calibrated vaporizer) and secured in a stereotaxic frame (RWD Life Science). Bilateral hippocampal injections were performed as follows:

CA3 injections: AAV9::*hSyn-Syp-mCherry-eGFP* was injected into CA3 of *Nrbf2* KO and WT mice (coordinates relative to bregma: AP −2.0 mm, ML ±2.35 mm, DV −2.25 mm from skull surface).

CA1 injections: In Nrbf2 KO mice, the right CA1 (AP −2.0 mm, ML +1.5 mm, DV −1.5 mm) received either AAV9::*GfaABC1D-eGFP-P2A-Nrbf2-3XFlag*, AAV9::*GfaABC1D-eGFP-P2A-Rab7-3XFlag*, or AAV9::*Syn-eGFP-P2A-Nrbf2-3XFlag*, while the left CA1 (ML −1.5 mm) received matched control vectors (AAV9::*GfaABC1D-eGFP-3XFlag*, AAV9::*GfaABC1D-eGFP-3XFlag*, or AAV9::*Syn-eGFP-3XFlag*).

All viruses (0.5 μL/site) were delivered at 0.05 μL/min using a motorized injector (Hamilton syringe, 5μL). After suturing and topical antiseptic application, mice recovered on a 37°C heating pad for 1 h before returning to home cages.

### Western blot

Astrocyte cultures were washed twice with ice-cold PBS and incubated for 30 min in ice-cold RIPA lysis buffer. Lysates were centrifuged at 12,000 × g for 30 min at 4°C, and the supernatant was collected. The concentration of protein was measured with a BCA assay kit (Pierce). The lysates were then heated at 100°C for 10 minutes. Following that, the cellular extracts were subjected to 12% SDS-PAGE to separate the proteins. These separated proteins were then transferred onto PVDF membranes using a wet transfer tank, which was immersed in an ice bath and maintained at 0.2 A for 1 hour. Subsequently, the membranes were blocked for 1 hour in a TBS with Tween (TBST) buffer containing 5% (w/v) milk powder. Next, the membranes were exposed to specific antibodies in TBST with 5% (w/v) BSA and incubated overnight at 4°C. To detect the protein bands, an HRP substrate (GE healthcare, 45-002-401) was employed, following the manufacturer’s instructions for chemiluminescence detection in a dark room. The visualization and analysis of the Western blotting results were performed using software (Image Lab 5.1, Bio-Rad, Munich, Germany).

### Single-cell RNA Sequencing and Data Analysis

#### Sample preparation

Fresh tissues were immediately used to prepare their single-cell suspension. Briefly, the collected fresh tissue was washed by DPBS (PBS without calcium and magnesium) and carefully chopped into small pieces. Enzymatic digestion was performed with collagenase II (2 mg/mL), collagenase IV (2 mg/mL) and dispase (0.2 mg/mL) under 37 °C for 15 - 25 minutes. The digested solution was transferred to a centrifuge tube through a 40 μm stainless nylon mesh (Greiner Bio-OneGmbH, Germany). The filtrate was immediately centrifuged at 500 g for 5 min, and discarded the supernatant slowly. The cell sediment was resuspended and lysis with RBC Lysis buffer on ice for 5 min to remove red blood cells, and then washed with DPBS at 500 g for 5 min twice. Cell concentration and viability were determined on a Cellometer Auto 2000 (nexcelom, USA) after AO/PI staining.

#### Fluorescence - activated cell sorting (FACS)

Immediately after dissociation, cells were resuspended in FACS buffer (5% FBS in PBS) and Fc-blocked for 10 min. This was followed by a wash and staining with anti-CD45 PE (BioLegend; San Diego, CA) for 30 min in sorting buffer (0.1% BSA in PBS) at 4 °C. After washing, viable CD45+ cells were sorted using the Beckman Coulter MoFlo Astrios. Subsequently, the cells were washed twice and re-suspended in a sorting buffer. A cell number and viability count were performed on a Cellometer Auto 2000 using the ViaStain™ AOPI Staining Solution (Nexcelom Bioscience LLC, Lawrence, MA, USA) immediately prior to scRNAseq.

#### 10x Genomics Chromium library construction and sequencing

The library synthesis was conducted in accordance with the instructions of Chromium Next GEM Single Cell 3ʹ Reagent Kits v3.1 (10× GENOMICS). In brief, the cells of each sample was adjusted to 1000 cells/μL. Cell suspensions were loaded on the Chromium Controller instrument to generate single-cell Gel Beads-In-Emulsions (GEMs), and individual cells were isolated into droplets together with gel beads that coated with unique primers bearing 10× cell barcodes, unique molecular identifiers (UMI), and poly (dT) sequences. In GEMs, individual cells were barcode-labeled and mixed with reverse transcriptase, and the GEM-reverse transcriptions were performed on a Veriti 96-well thermal cycler (Thermo Fisher Scientific, Waltham, MA, USA). Then the cDNA libraries were amplified, fragmented, end repaired, and A-tailed after reverse transcription; then adaptor was ligated after size selection; finally, sample index PCR was performed and SPRI select beads were used for final purification. At last, Libraries were sequenced using the NovaSeq 6000 platform (Illumina) to a depth of approximately 500 million reads per library with 2×150 read length.

#### scRNA - seq data analysis

Raw snRNA-seq data were analyzed using Seurat v4.4.0^26^. Initial quality control involved the removal of cells with UMI counts less than 200 or greater than 6000, as well as cells exhibiting >5% expression of mitochondrial, ribosomal, or erythrocyte-associated genes. DecountX^27^ and DoubletFinder^28^ were applied using default parameters to eliminate low-quality cells further to minimize the impact of ambient RNA contamination and potential doublets. The resulting data were scaled using the *SCTransform* procedure, and batch effects were corrected using *Harmony*. Major cell type clusters were identified at a resolution of 0.1, and Uniform Manifold Approximation and Projection (UMAP) was used for dimensionality reduction and visualization of cell distributions. Cell type annotation was performed based on the expression of canonical marker genes: Astrocytes (*Aldh1l1, Aqp4, Gpc5, Gja1, Gfap*), Microglia (*C1qa, Cd74, Ccl3, Csf1r, P2ry12, Tmem119, Cx3cr1, Hexb*), Excitatory neurons (*Cbln2, Ldb2, Nrgn, Slc17a7, Neurod6*), Inhibitory neurons (*Gad1, Gad2, Slc32a1*), Endothelial cells (*Pecam1, Cdh5, Ebf1, Flt1, Cldn5*), and Oligodendrocytes (*Mbp, Plp1, Oligo1*). We used a resolution of 0.3 to identify astrocyte subclusters. We used the FindAllMarkers function and the MAST method to identify differentially expressed genes (DEGs) specific to astrocyte subclusters and corrected for sex (parameters: test.use = ‘MAST’, latent.vars = ‘Sex’). The *scProportionTest*^29^ was used to assess the distribution of astrocyte subclusters across different experimental groups, with Benjamini-Hochberg correction applied for multiple testing. Gene ontology enrichment analysis was performed using Metascape on the top 20 DEGs for each astrocyte subcluster. To investigate the potential relationship between astrocyte subclusters and aging, an aging-associated gene set was obtained from the GenAge database(https://genomics.senescence.info/genes). An aging score for each astrocyte subcluster was calculated using the *AddModuleScore* function. All analyses were performed in R (version 4.3.1). Two-group comparisons were performed using a two-tailed Student’s t-test, while comparisons involving more than two groups were performed using analysis of variance (ANOVA) followed by Benjamini-Hochberg correction. A P-value of <0.05 was considered statistically significant.

### Spatial Transcriptomics and Data Analysis

#### ST tissue handling for OCT

Tissues were rapidly frozen in optimal cutting temperature (OCT) compound (4583, Sakura) and stored at −80°C. Cryosections of 10-μm thickness were prepared using a cryostat (CM3050S, Leica) at −20°C. The optimal permeabilization time for tissue sections was determined to be 15 minutes following the manufacturer’s instructions (10x Genomics, Visium Spatial Tissue Optimization). Sections were mounted on Visium Spatial Gene Expression Slides (PN-1000184, 10X Genomics) and fixed in pre - cooled methanol (32213, Sigma-Aldrich) for 5 minutes. Hematoxylin (51275, Sigma-Aldrich) and eosin (HT110116, Sigma-Aldrich) staining were performed for histological annotation. Spatial gene expression libraries were then generated according to the instructions of the Visum Spatial Gene Expression Kit from 10X Genomics. The libraries were sequenced using the Illumina NovaSeq6000 platform to generate approximately 150 M read-pairs per section.

#### ST tissue handling for FFPE

The tissues were obtained from patients after resection, fixed by formalin, and embedded in paraffin. RNA quality assessment of FFPE tissue blocks was performed by calculating the percentage of total RNA fragments > 200 nucleotides (DV200) of RNA extracted from tissue sections. For the high RNA quality, DV200 > 30% was required. A 10x FFPE gene expression slide (PN-1000185,10X Genomics) was used for ST. A slide with 5 - μm FFPE section was dewaxed with xylene (214736, Sigma-Aldrich) and stained with hematoxylin (51275, Sigma-Aldrich) and eosin (HT110116, Sigma-Aldrich). After visualizing and scanning the whole slide, decrosslinking was performed using the tris-thylenediaminetetraacetic acid (TE) buffer (10-0046, GeneMed) to release the RNA. Forward and reverse human transcriptome probes (PN-1000364, 10X Genomics) were used for probe hybridization overnight. After hybridization, the cDNA libraries were constructed according to the 10X protocol (CGO00407_VisiumSpatialGeneExpressionforFFPE_UserGuide_RevA). The sequencing was performed on NovaSeq 6000 (Illumina) platform to generate approximately 150 M read-pairs per section.

#### Spatial RNA - seq data processing, visualization and Analysis

Raw spatial transcriptomics data were analyzed using Giotto v3.1^30^. Initial quality control was performed using the *filterGiotto* function with the following parameters: *expression_threshold* = 0.5, *feat_det_in_min_cells* = 50, and *min_det_feats_per_cell* = 1000. Highly variable genes (HVGs) were identified for data scaling and dimensionality reduction. Spatially-dependent subclusters were identified using the *doLeidenCluster* function. Given that each spot contains expression profiles from multiple cells, we used *FindTransferAnchors* to transfer cell type labels from the snRNA-seq data to the spatial transcriptomics data, outputting a probabilistic classification of each spot to each snRNA-seq derived cell type. To investigate the transcriptomic features of SAD spots, given that SAD was labelled with green fluorescence, fluorescence images were registered to the spatial transcriptomics images using ImageJ, and the green channel was isolated for quantification. Subsequently, fluorescence images were segmented using QuPath, with spot locations derived from the *tissue_position* data. Spot parameters were set to *tissue_hires_scalef* = 0.13832216 and *spot_diameter_fullres* = 139.30696255752676. The segmentation images generated by QuPath were transferred to ImageJ, and the *multi-measure* function was used to quantify green fluorescence intensity for each spot. Integrating spot fluorescence intensity with spatial transcriptomics data, we used *FindMarkers* to identify differentially expressed genes (DEGs) between SAD spots and their immediately adjacent spots.

### Primary astrocyte cultures

Primary astrocytes were isolated from neonatal mice (P1, sex not distinguished). Pups were terminally anesthetized with isoflurane and decapitated. Whole brains were rapidly dissected into ice-cold HBSS buffer containing 2% penicillin-streptomycin. Hippocampi and cortices were microdissected under a stereomicroscope, digested with 0.125% trypsin-EDTA for 15 min at 37°C, and dissociated. The cell suspension was filtered through a 100 μm strainer and plated on flasks. Cultures were maintained in DMEM/F12 with 20% FBS, 100 U/ml penicillin, and 100 μg/ml streptomycin at 37°C/5% CO_2_. After 7 days, microglia and oligodendrocyte precursors were removed by orbital shaking (250 rpm, 37°C, 30 min), pure astrocytes as verified by GFAP immunofluorescence.

### Primary astrocyte transfection

Primary astrocytes were harvested as described previously and plated in six-well plates. AAV9::*GfaABC1D-eGFP-P2A-Nrbf2-3XFlag*, AAV9::*GfaABC1D-eGFP-P2A-Rab7-3XFlag*, or AAV9::*Syn-eGFP-P2A-Nrbf2-3XFlag* was diluted in 1 mL of DMEM/F12 medium per well at an MOI = 10 and infected astrocytes for 4 hours, followed by supplementation with 1 mL of complete medium (DMEM/F12:FBS = 10:1) per well. Repeat infection after 72 hours. Astrocytes were transfected at DIV 8–10.

### Synaptosome Purification

To isolate subcellular fractions from brain tissue, olfactory bulbs and cerebellum were carefully excised while preserving the cerebral cortex and hippocampus. The dissected tissues were then rinsed multiple times with ice-cold PBS to remove blood and debris. Half of the combined cortical and hippocampal tissues were transferred to a microcentrifuge tube containing 500 μL of ice-cold sucrose buffer (0.3 M sucrose, 20mM HEPES 7.5, 1X protease inhibitor tablet, 2 mM DTT, 0.5X phosphotase inhibitor cocktail). All subsequent steps were performed at 4°C or on ice until the addition of SDS. Tissues were homogenized using a tight-fitting dounce homogenizer with 6–10 gentle up-and-down strokes. The homogenate was rotated in a cold room for 20 min to ensure complete lysis, then centrifuged at 300×g for 3 min in a 600 μL tube. The supernatant was transferred to a fresh tube and centrifuged at 10,000×g for 10 min. The resulting pellet, representing the crude synaptosome fraction, was resuspended in 0.5 mL of sucrose buffer, centrifuged again at 10,000×g for 10 min, and the supernatant was discarded. The washed synaptosome pellet was resuspended in 500 μL of buffer B (20 mM Tris 7.5, 1.5% SDS, 1mM DTT), boiled for 5 min to denature proteins, and stored at −80°C for following phagocytosis experiment. For synaptosome electron microscopy, the synaptosome pellet was fixed by 500uL Electron Microscope Fixative at RT for 2 h, keep 4℃ for delivery and TEM.

### Synaptosomes conjugation to pHrodo

pHrodo Red, succinimidyl ester (P36600) and were dissolved as described in the manual. Synaptosome samples in 1.5-mL microcentrifuge tubes were pelleted by centrifugation at 15000 × g for 3–4 min at 4°C. The supernatant was carefully aspirated, and pHrodo dye was added at a ratio of 1 μL per 0.3 mg of synaptosome protein. Tubes were wrapped in aluminum foil to protect from light and incubated on a twist shaker at 30–40 rpm for 30 min at room temperature. Following incubation, 1 mL of PBS was added to each tube to halt the reaction. Samples were centrifuged again at 15000 × g for 1–2 min, and the supernatant was discarded. The synaptosome pellet was gently resuspended in 1 mL of PBS by pipetting and washed three additional times via centrifugation and resuspension to remove unbound dye. After the final wash, the pellet was resuspended in 500 μL of DMEM/F12 medium for immediate use or storage at −80°C.

### Phagocytosis Live-Imaging Assay

Primary astrocytes were seeded in 96-well imaging plates at a density optimized for confluency and cultured for 2 days to allow stable adherence. Prior to imaging, culture media were aspirated, and cells were washed twice with 100 μL of pre-warmed PBS per well. Wells were then supplemented with 100 μL of complete medium containing pHrodo-conjugated synaptosomes. Live imaging was performed using the Opera Phenix Plus High-Content Screening System with a 40x water immersion objective. The system was pre-warmed to 37°C with 5% CO_2_ humidity. A total of 50 time points were acquired over 4 hours at 5-minute intervals, covering ⩾9 fields per well to ensure statistical robustness. Image analysis was conducted using Harmony High-Content Imaging and Analysis Software. Phagocytic activity was quantified by calculating the number of internalized synaptosomes per cell, mean fluorescence intensity of intracellular pHrodo, and the percentage of phagocytic cells.

### Morris water maze

The Morris water maze test was performed in accordance with established protocols to evaluate spatial memory performance^31^. The device is a circular white water pool (120 cm diameter × 50 cm depth) filled with water dyed white with TiO2 at a constant temperature of 22 °C. A 10-cm-diameter platform was submerged 1 cm below the water surface at a fixed position. Different shapes and colors were pasted on the side walls of the swimming pool as visual stimuli to train the mice to navigate and locate the hidden platforms in the pool. One day before the hidden platform training, mice underwent three trials of pre-training by swimming through a channel to mount a rescue platform. During all hidden platform training trials, the platform was kept in the same submerged position; the position at which the mice fell into the pool was different between trails. For the hidden trials, mice were subjected to three trials per day for five days. Each trial lasted 60 s or until the mouse found the platform. If the mouse did not find the platform during the allocated time, the experimenter directed the mouse to the platform and placed on the platform for 10 s. On the 6th day, the platform was removed for a probe trial (60 s) to assess long-term spatial memory retrieval. All parameters were recorded by a video tracking system (Labmaze V3.0, Zhongshi Technology).

### Contextual fear conditioning

The contextual fear conditioning was conducted in Startle and Fear Combined System (Panlab, Harvard Apparatus). This chamber was a 27 cm × 27 cm arena with a conductive metal grid floor. The fan provides 60 dB of background white noise.

For training day, mice were exposed to an 180s period for free exploration followed by three consecutive 2s foot shocks of 0.5 mA at 100 s intervals and a final 60s resting period. On the testing day, mice were re-exposed to the same chamber for 180s. Freezing behavior was defined as absent of movement other than breathing for at least 2 s. Percentage time freezing was scored by an automated motion-sensitive software for behavior boxes (PACKWIN V2.0, Harvard Apparatus).

### Y maze

The Y-maze spontaneous alternation performance (SAP) test assesses an animal’s capacity to recognize and differentiate a previously explored environment. This apparatus comprises three arms, each measuring 30 cm in length, 20 cm in height, and 6 cm in width, arranged at 120° angles relative to one another. Mice were introduced at the central junction of the maze and permitted to explore freely for a duration of 10 minutes. To prevent olfactory cues from influencing behavior, the maze was disinfected with a 75% ethanol solution between trials. Arm entries were recorded using the Labmaze video tracking system (Labmaze V3.0, Zhongshi Technology). SAP was determined by counting the number of consecutive entries into a novel arm within a sequence of three choices. The spontaneous alternation percentage was calculated as follows: (number of actual alternations / [total arm entries – 2]) × 100.

### LTP

Mature adult mice were deeply anesthetized via intraperitoneal injection of Avertin (250 mg/kg; 0.2 mL of 1.25% Avertin working stock solution per 10 g body weight) and transcardially perfused with 25–30 mL of ice-cold carbogenated (95% O_2_/5% CO_2_) choline chloride aCSF (120mM Choline chloride, 204mM KCL, 7mM MgCl2·6H2O, 0.5mM CaCl2·2H2O, 1.25mM NaH2PO4·H20, 5mM Sodium ascorbate, 3mM Sodium pyruvate, 26mM NaHCO3, 25mM D-glucose). Following perfusion, mice were decapitated, and brains were rapidly extracted (<1 minute) and submerged in ice-cold choline chloride aCSF for an additional minute. Transverse hippocampal slices (350 μm) were prepared using a vibratome (Campden Instruments, UK) in ice-mash choline chloride aCSF continuously gassed with 1 95% O_2_/5% CO_2_.

Slices were transferred to a recovery chamber containing oxygenated standard aCSF (10 mM D-glucose, 3.5 mM KCl, 1.25 mM KH2PO4, 2.5 mM CaCl2, 1.5 mM MgSO4, 120 mM NaCl, and 26 mM NaHCO3) at 32°C for 1 hour, then maintained at room temperature in carbogenated aCSF until recording. For electrophysiological experiments, slices were placed in a MED Prob 16 recording chamber (Alpha MED Scientific, MED-PG5001A) equipped with a 4×4 grid of planar microelectrodes. The CA1/CA2 stratum radiatum was positioned over the electrodes to enable stimulation of Schaffer collaterals and recording of field excitatory postsynaptic potentials (fEPSPs) from CA1 dendritic layers.

Extracellular recordings were performed using the MED64-Quad II system with Mobius Pro software (Alpha MED Scientific). Biphasic current pulses (10–60 μA, 0.2 ms duration) were delivered every 20 seconds to evoke fEPSPs. Stimulus intensity was calibrated for each slice using input/output (I/O) curves, with the test stimulus set to evoke 40% of the maximum fEPSP amplitude. After establishing a stable 20-minute baseline, long-term potentiation (LTP) was induced using theta burst stimulation (TBS): 5 trains of 10 bursts at 100 Hz, with 200 ms inter-burst intervals and 30 s inter-train intervals. fEPSP slopes were continuously monitored for 60 minutes post-TBS to assess LTP maintenance. Data were analyzed offline using custom scripts in MATLAB (MathWorks) to quantify changes in fEPSP slope relative to baseline values.

### GST-R7BD Affinity Isolation

GST-R7BD (GST-RILP) was purified from bacterial cultures as previously described^16^. Briefly, the GST-R7BD construct was transformed into Escherichia coli strain BL21 (Takara, 9126). A 300 mL 2XYT culture was incubated at 30°C for 12 h. Bacteria were pelleted by centrifugation at 5,000 × g for 10 min at 4°C, washed with cold phosphate-buffered saline (PBS), and resuspended in 8 mL of cold lysis buffer (50 mM Tris-HCl, pH 7.5, 1 mM EGTA, 1% Triton X-100, 0.27 M sucrose, 0.1% β-mercaptoethanol (Sigma, M6250), 1:4,000 DNase I (Thermo Fisher, EN0521), and protease inhibitors). Protein purification was performed using 500 μL of glutathione-Sepharose 4B beads (GE Healthcare, GE17-0756-01). pGEX-4T-3-mR7BD was a gift from Aimee Edinger (Addgene plasmid # 79149; http://n2t.net/addgene:79149; RRID: Addgene_79149)

Mammalian cells were lysed in pull-down buffer (20 mM HEPES, 100 mM NaCl, 5 mM MgCl_2_, 1% Triton X-100, protease inhibitors). For each affinity isolation, 500 μg of cell lysate was incubated with 100 μL of pre-equilibrated beads in 1 mL of affinity-isolation buffer overnight at 4°C with gentle rocking. Beads were washed twice with cold affinity-isolation buffer, and bound proteins were eluted by adding 1× sample buffer and incubating at 95°C for 8 min. The amount of GTP-bound RAB7 was determined by immunoblotting using a RAB7-specific primary antibody and appropriate secondary antibody.

### Time-lapse microscopy and analysis

Live imaging was conducted using the Opera Phenix Plus High-Content Screening System with corresponding acquisition software. The cells were maintained in an environmental chamber at 37°C with 5% CO_2_ humidity throughout the process. For the experiments, primary astrocytes were first pre-stained with LysoTracker Green for 30 minutes at 37°C to label lysosomes, followed by washing with pre-warmed HBSS to remove unbound dye. LysoTracker-stained lysosomes were performed using a 40× water immersion objective. Images were acquired every 5 minutes for up to 4 hours to monitor lysosomal trafficking.

For Raichu-Rab7 expression, primary astrocytes were transfected with the respective plasmids using a lipo3000 according to the manufacturer’s instructions. Forty-eight hours post-transfection, cells were treated with unstained synaptosome. Live-cell imaging of Rab7 dynamics was performed using a 63× water immersion objective. Images were acquired every 20 minutes for up to 40 minutes to monitor Rab7 recruitment to autophagic structures. FRET efficiency analysis for pRaichu-Rab7 was conducted using the Opera system’s built-in FRET module, which quantitated donor (488 nm excitation, 510-530 nm emission) and acceptor (561 nm excitation, 580-600 nm emission) signals. The FRET ratio (acceptor/donor intensity) was calculated pixel-wise across the cell population to assess Rab7 conformational changes or interactions, with background subtraction applied using non-transfected control cells. Data were exported and analyzed using Harmony High-Content Imaging and Analysis Software for statistical comparisons between groups.

### Reagents

Antibody
| name | Used for | Manufacturer and Catalog Number |
| --- | --- | --- |
| p62 | IF and IHC | Abcam, ab109012 |
| p62 | IF and IHC | Progen, GP62-C |
| GFAP | IF | Cell Signaling Technology, 12389 |
| GFAP | IF and WB | Merck,G3893 |
| Synaptophysin | IF | Abcam, ab32594 |
| PSD95 | IF | CST, 3450 |
| Iba1 | IF | FUJIFILM Wako, CAJ3125 |
| CD31 | IF | Abcam, ab64543 |
| β3Tubulin | IF | Cell Signaling Technology, 5568 |
| NeuN | IF | Millipore, MAB377 |
| CNPase | IF | Cell Signaling Technology, 5664 |
| LAMP1 | IF and WB | Abcam, ab24170 |
| cathepsin B | WB | Cell Signaling Technology, 31718 |
| cathepsin D | WB | Santa Cruz, sc-377299 |

## Data availability

The datasets generated during and/or analyzed during the current study are available from the corresponding author on reasonable request. Upon acceptance of the manuscript, we will upload the data as required. All data related to this study are welcome to be requested from the authors.

## Author contributions

J. Lu conceived the study and designed the experiments. C. Ming and S. Wang performed all omics data analysis. E. Wang, Y. Wang, X. Zhuang, W. Wang, and M. Wu conducted the animal experiments. E. Wang, Y. Wang, and L. Xie were responsible for sample processing and other molecular biological experiments. Y. Wang, K. Cheung, and A. Wu carried out the electrophysiological and behavioral experiments. E. Wang wrote the manuscript, with contributions from K. Yang, Z. Song, J. Li, C. Liu, L. Wang, Y. Yao, M. Li, H. Shen, H. Su, and Z. Yue. K. Yang performed electron microscopy experiments and prepared the relevant figures. Z. Song organized the images and data. J. Li conducted protein purification and other molecular biological experiments. C. Liu and L. Wang were in charge of transgenic animal construction and breeding. Y. Yao processed the macaque brain samples. All authors discussed the results and commented on the manuscript.

